# Intestinal cell kinase regulates chondrocyte proliferation and maturation during skeletal development

**DOI:** 10.1101/139089

**Authors:** Mengmeng Ding, Li Jin, Lin Xie, So Hyun Park, Yixin Tong, Di Wu, Zheng Fu, Xudong Li

## Abstract

An autosomal recessive loss-of-function mutation R272Q in human *ICK* (intestinal cell kinase) gene causes profound multiplex developmental defects in human ECO (endocrine-cerebro-osteodysplasia) syndrome. ECO patients exhibit a wide variety of skeletal abnormalities, yet the underlying cellular and molecular mechanisms by which ICK regulates skeletal development remain largely unknown. The goal of this study is to understand the structural and mechanistic basis underlying skeletal anomalies caused by ICK dysfunction. *Ick* R272Q knock in transgenic mouse model not only recapitulated major ECO skeletal defects such as short limbs and polydactyly but also revealed a deformed spine with deficient intervertebral disc. Loss of ICK functions markedly reduces mineralization in the spinal column, ribs, and long bones. *Ick* mutants show a significant decrease in the number of proliferating chondrocytes and type X collagen-expressing hypertrophic chondrocytes in the spinal column and the growth plate of long bones. Our results demonstrate that ICK plays an important role in bone and intervertebral disc development by promoting chondrocyte proliferation and maturation, and thus provide novel mechanistic insights into the skeletal phenotypes of human ECO syndrome.

## Introduction

Endochondral ossification is a stringently controlled and coordinated process during axial bone development, starting from mesenchymal stem cell condensation to form a template of future bone. Chondrocytes at the end of the template proliferate, while chondrocytes in the middle of the template differentiate into hypertrophic chondrocytes that secrete extracellular matrix (ECM). Hypertrophic chondrocytes eventually undergo apoptosis, and leave the cartilaginous matrix scaffold that attracts osteoblasts to form the primary spongiosa [1–2].

Primary cilium, a microtubule-based organelle, has been implicated an important role in the skeletal development and homeostasis [3]. Mutations interfering with formation and maintenance of primary cilia were identified in skeletal ciliopathies, including Ellis-van Creveld syndrome (EVC), cranioectodermal dysplasia (CED), asphyxiating thoracic dystrophy (ATD), short rib-polydactyly syndrome (SRPS), and endocrine-cerebro-osteodysplasia (ECO) syndrome [4]. ICK, a highly conserved and ubiquitously expressed serine/threonine protein kinase in human kinome, plays an important role in the maintenance of primary cilium. Several loss-of-function point mutations in human *ICK* gene, including c.1305G-A (R272Q), c.358G-T (G120C), and c.238G-A (E80K), were reported in ECO and ECO-like syndromes that display profound skeletal abnormalities such as polydactyly, short ribs, bowed limbs, and abnormal long bones [5–9].

Our prior work has demonstrated an important role for ICK in the regulation of cell proliferation and survival *in vitro* [10–11]. An essential role of ICK *in vivo* emerged from the report of human ECO and ECO-like syndromes [5–7]. Recently, *Ick* knockout mouse models reproduced ECO phenotypes in the cerebral and skeletal systems and linked ICK deficiency to abnormal structures of primary cilia [7–9]. However, cellular and molecular mechanisms of ICK in skeletal phenotypes are still elusive. In order to advance our understanding of the structural and mechanistic basis underlying skeletal anomalies caused by ICK dysfunction, we generated an *Ick* R272Q knock-in mouse model that can recapitulate ECO developmental phenotypes including skeletal defects. We analyzed the fetal bones of *Ick* R272Q mutant embryos with respect to skeletal phenotype, chondrocyte proliferation and differentiation. Our results demonstrate that ICK plays a critical role in the development of axial skeleton and intervertebral disc (IVD) by regulating chondrocyte proliferation and maturation.

## Materials and Methods

### Generation of *Ick*^R272Q^ knock-in mutant mice

All procedures involving animals were performed in accordance with ethical standards in an animal protocol that was approved by the Institutional Animal Care and Use Committee. The R272Q (CGA>CAA) point mutation was introduced into the exon 8 of the wild-type (WT) *Ick* allele on a bacterial artificial chromosome (BAC) to generate *Ick*/R272Q BAC. A LNL (LoxP-Neo-LoxP) cassette was inserted in the intron downstream of exon 8. A gene targeting vector was constructed by retrieving the 5kb long homology arm (5’ to LNL), the LNL cassette, and the 2kb short homology arm (3’ to LNL) into a plasmid vector carrying the DTA (diphtheria toxin alpha chain) negative selection marker. The LNL cassette conferred G418 resistance during gene targeting in PTL1 (129B6 hybrid) ES cells and the DTA cassette provided an autonomous negative selection to reduce the random integration event during gene targeting. Several targeted ES cell clones were identified and injected into C57BL/6 blastocysts to generate chimeric mice. Male chimeras were bred to homozygous EIIa (cre/cre) females (in C57BL/6J background) to excise the neo cassette and to transmit the *Ick*/R272Q allele through germline (Precision Targeting Lab, USA). *Ick^R272Q^* heterozygous were normal and inter-crossed to produce *Ick^R272Q^* homozygous embryos. Animals were housed in a temperature controlled colony room on a 12-hour light cycle, and had access to food and water ad libitum. For timed pregnancy, the presence of a copulation plug in the morning represented embryonic day (E) 0.5. Pregnant mice were euthanized by CO_2_ inhalation and embryos were harvested at different developmental time points (n=2–5).

### Whole-mount skeletal staining

Embryos at E15.5 or E18.5 were placed in PBS after euthanization. The skin, internal organs and bubbles were removed from the body cavity, and then fixed in 95% Ethanol. The cartilage was stained with Alcian Blue and bone was stained with Alizarin Red following a standard protocol [12]. After clearing, samples were placed in 100% glycerol for long-term storage and photography (Olympus SZX12, Olympus).

### Alcian blue and Picrosirius red staining

Embryos were washed with PBS and fixed in 100% ethanol overnight. Paraffin embedded sections (5µm) were subjected to 1% Alcian blue solution (pH 2.5) and Picrosirius red (0.1% Sirius red in saturated aqueous picric acid) staining. Images were taken with a Nikon Eclipse E600 microscope. Five semi-serial sections (10 µm between each level) through the center of femurs and spine from each mouse were cut, stained, and photographed for measurement using an NIS element BR imaging software (Nikon, Japan).

### Von kossa staining

Paraffin embedded sections were dewaxed and incubated in 5% silver nitrate solution (w/v, Sigma, MO) under UV light for 30 minutes. Sections were rinsed in distilled water and immersed in 5% sodium thiosulfate (Sigma, MO) for 5 mins and counterstained with hematoxylin (Sigma) for 5 mins.

### Immunohistochemistry

Immunostaining for paraffin sections was performed as described previously [13–14]. For collagen X (GTX105788, 1:400, GeneTex, CA) and collagen II (CB11,1:100, Chondrex, WA) staining, sections were treated with 2% bovine testicular hyaluronidase 30 minutes for antigen retrieval. For phospho-H3 (H0412,1:200, Sigma, MO) staining, sections were immersed in 10 µM of sodium citrate solution at 85°C for 30 minutes for antigen retrieval. The standard 3,3'-diaminobenzidine (DAB) procedure was followed to visualize the signal. Hematoxylin was used for counter-staining. IgG control was performed following identical procedures excluding the primary antibody.

### Micro-computed tomography (micro-CT)

Embryos at 18.5 were fixed in 10% formalin. Micro-CT scans were performed on a micro-CT 80 scanner (70 kV, 114 μA; Scanco Medical, Switzerland), and then were reconstructed with an isotropic voxel size of 10 μm. Multi-level thresholds procedure (threshold for bone = 225) was applied to discriminate soft tissue from bone. Three-dimensional images were acquired for qualitative evaluation in an X-ray image mode.

### Statistical analysis

Histological quantification data were expressed as mean±SD. Statistical comparison between genotypes was performed by a two-tailed student’s *t*-test. *p*<0.05 was considered significant.

## Results

### Overall morphological defects in *Ick*^R272Q^ homozygous mutant embryos

Previously we have shown that ICK is ubiquitously expressed in mouse tissues [15]. We first examined ICK expression in bone and cartilage tissues by Western blotting and by β-galactosidase staining of an *Ick-LacZ* reporter mouse. As shown in supplementary Fig.1A, ICK protein was detected in tibia, sternum, and rib at a level that is comparable to that of lung and heart. A truncated *Ick* transcript fused with *LacZ* was expressed in *Ick^tm1a/+^* heterozygotes (from NIH Knockout Mouse Project Repository) [9], thus LacZ staining can be used to map ICK expression in mouse tissues. In supplementary Fig.1B, LacZ staining of E15.5 whole embryos clearly indicates ICK expression in limbs and spine.

**Figure 1:**
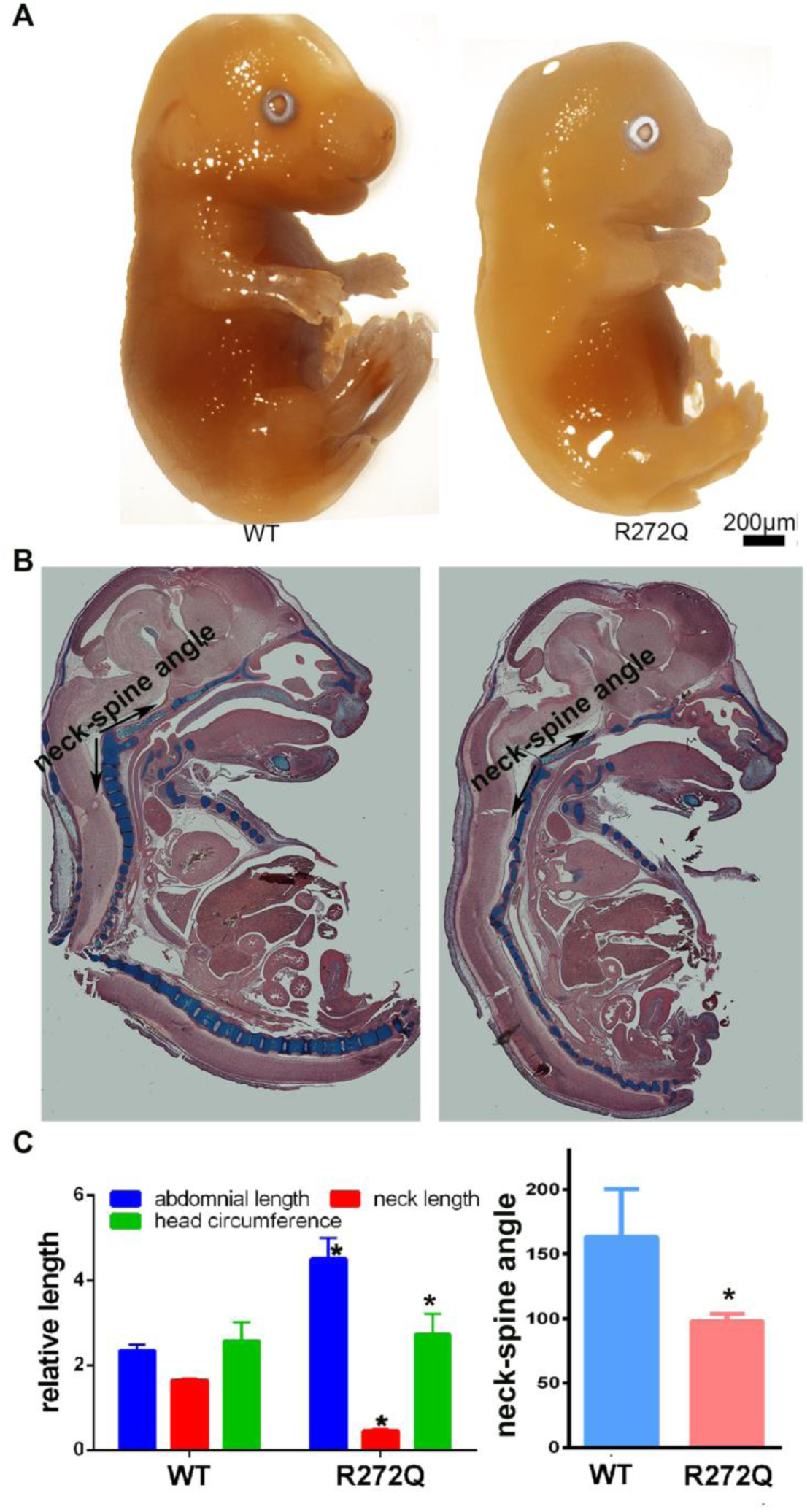
Gross morphological abnormalities of *Ick* R272Q homozygous mutant. (**A**) Sagittal views of *Ick* mutant (R272Q) and wild type (WT) littermate at E15.5. (**B**) Alcian blue staining showing distorted spine in *Ick* mutant at E15.5 as compared with WT littermate. (**C**). Quantification data indicating significantly increased abdominal length, decreased neck length, head circumference, and neck-spine angle in *Ick* mutant embryos (n=3).

Consistent with *Ick* R272Q being an autosomal recessive mutation in ECO syndrome, *Ick*^R272Q/+^ heterozygous mice were phenotypically indistinguishable from WT littermates. *Ick*^R272Q/+^ heterozygotes were interbred to generate *Ick*^R272Q/R272Q^ homozygotes that succumbed to death shortly after birth. Embryos from *Ick*^R272Q/+^ interbreed were harvested at E15.5 and E18.5. Gross morphology of *Ick*^R272Q/R272Q^ mutants is remarkably distinct from that of WT or heterozygous mutant littermates. *Ick* R272Q homozygous mutants have a broader head, smaller nose, bigger mouth, and shorter thorax. They also display broader cervical flexure, shorter distance between lower jaw to the forelimb, and larger abdominal cavity, flat ankles, and flat neck-spine angles (Fig. 1). These features closely resemble the clinical manifestations of ECO syndrome [9]. In addition to reported ECO skeletal phenotypes, these mutant mice also exhibit severe defects in the spinal column (Fig. 1b).

### Skeletal defects in *Ick*^R272Q^ homozygous mutant

To further investigate the role of ICK in bone and cartilage, whole mount skeletal analysis was performed on *Ick* mutant embryos. The skeletal phenotypes of *Ick*^R272Q/R272Q^ include short limbs, polydactyly, bowed long bones and much less bone mineralization (Fig. 2A). At E15.5, limb and tarsal bones of *Ick* mutants are all deviated; longitudinal length of long bones is significantly reduced, and polydactyly is evident in both forelimb and hindlimb (average 6.5 digits) as compared with WT littermates. A striking new discovery made in our *Ick* mutant mouse model that was not previously reported is the dramatic deformation of the spine. Compared with WT, the vertebrae and transverse processes of *Ick* mutants are severely deficient as shown by whole mount skeletal staining and micro-CT at E18.5 (Fig. 2B). Impaired mineralization of rib cage may not support the initial breath after birth.

**Figure 2:**
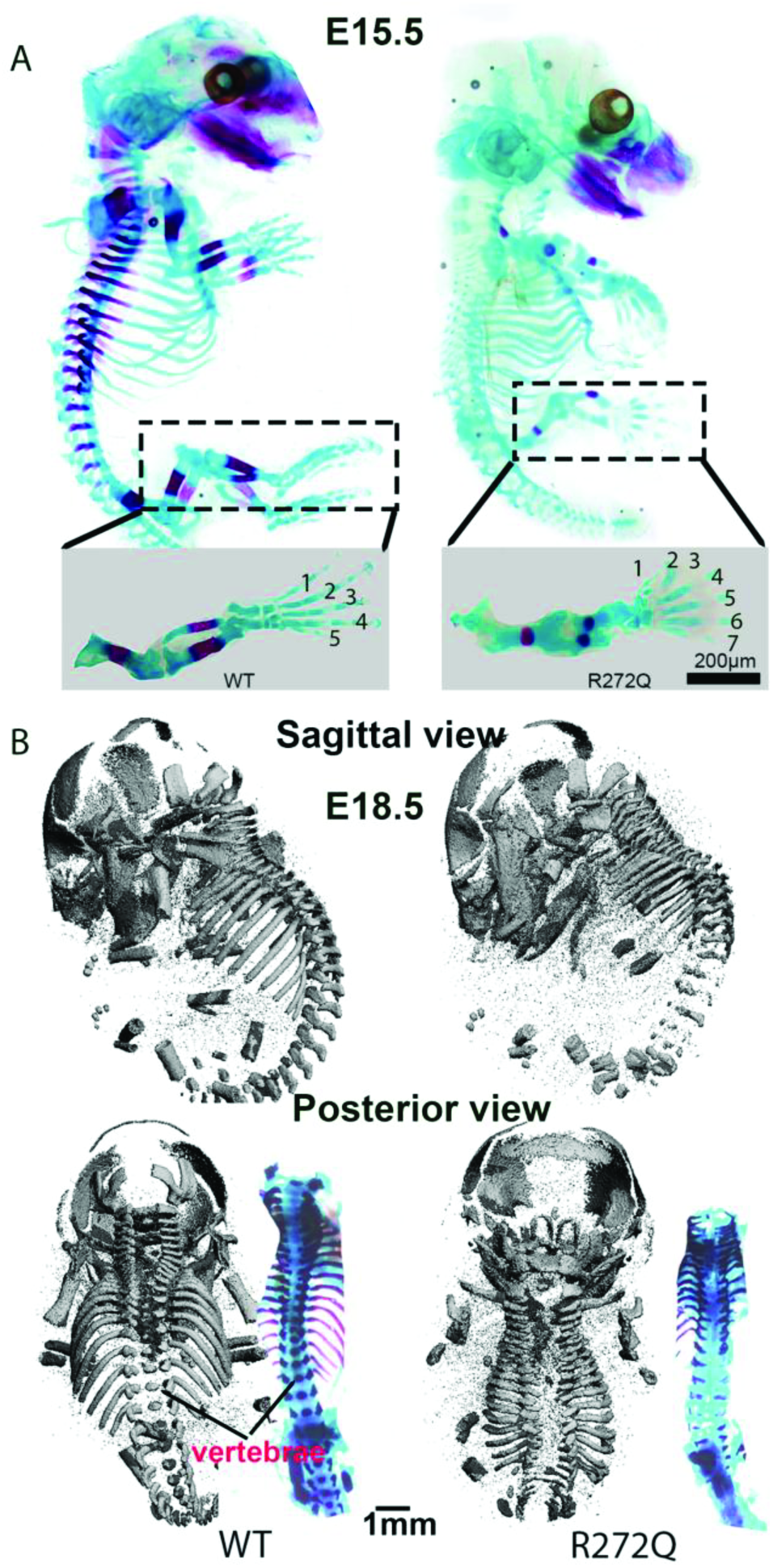
Skeletal defects of *Ick* R272Q homozygous mutant. (**A**) Whole mount skeletal staining of E15.5 embryos of *Ick* mutant and WT using Alizarin Red S (calcified tissue in red) and Alcian blue (cartilage tissue in blue). Note, *Ick* R272Q embryos show polydactyly, reduced mineralization (red), as well as bowed long bones. (**B**) Micro-CT images showing severe deficiency in vertebrae and spinal process of E18.5 *Ick* R272Q embryo.

### Abnormal phenotype of growth plate and intervertebral disc

Histological analyses were performed on femurs and spines of E15.5 and E18.5 embryos. *Ick* mutant femur is much shorter. While the size of cartilage region of *Ick* mutant femur is not significantly different from that of WT femur, endochondral bone is barely detectable at E15.5 femurs and at least two fold less at E18.5 femurs in *Ick* mutants as compared with controls, resulting in severely shortened bone (Fig. 3). The relative length of proliferating chondrocyte (PC) zone normalized to PC and resting chondrocyte (RC) zone is significantly shorter in *Ick* mutant than in WT littermates (Fig. 3). Von Kossa staining confirms much less matrix mineralization in *Ick* mutant femurs at E15.5 and E18.5 (Fig. 5).

**Figure 3:**
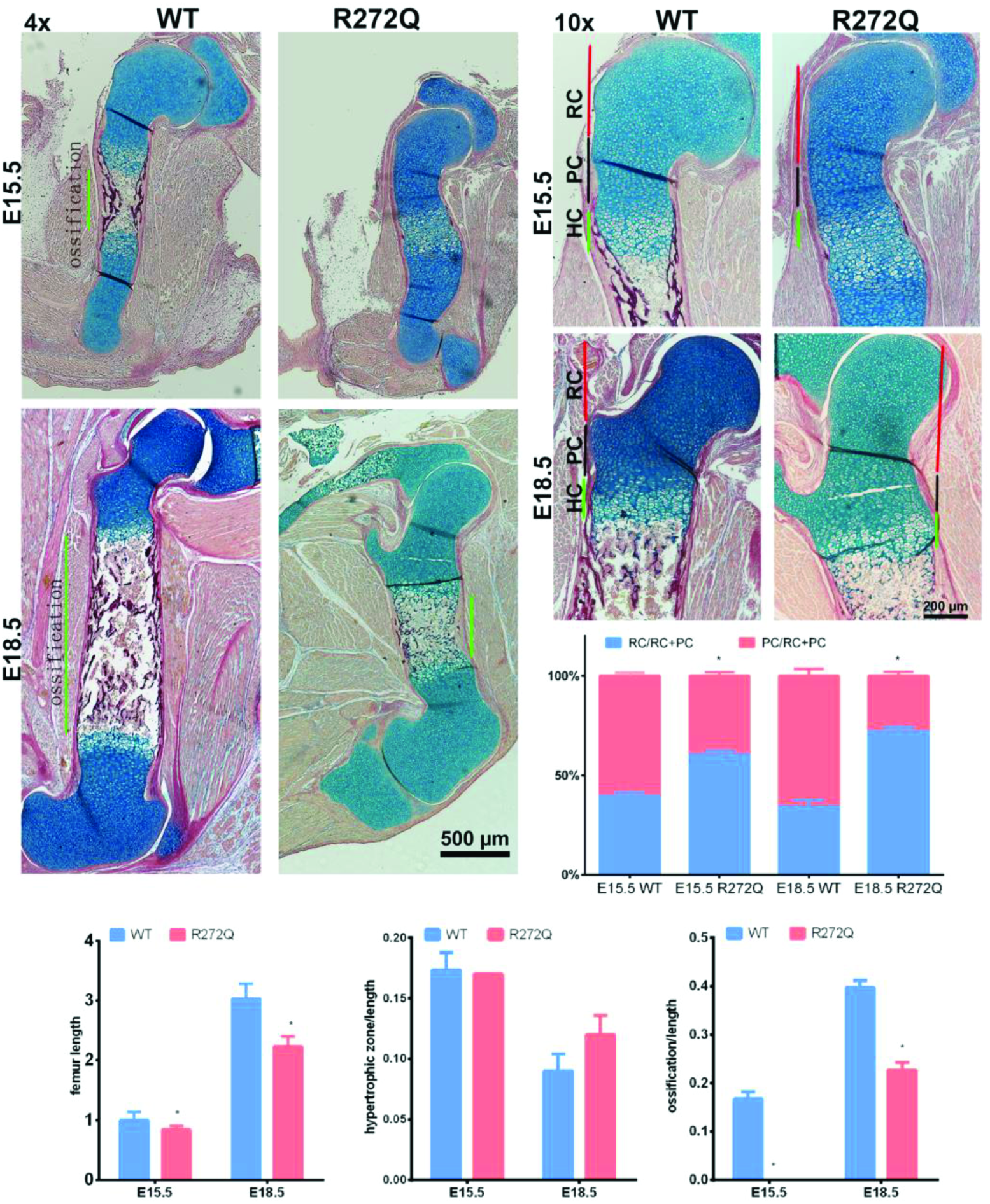
Bone ossification is significantly reduced in *Ick* R272Q mutant embryos. Paraffin embedded femoral sections were stained with Alcian blue and nuclear fast red. At E15.5, hypertrophic chondrocytes locate in the ossification center of *Ick* mutant embryos, whereas mineralized bone form in the WT littermates. At E18.5, bone ossification was observed in *Ick* mutant embryos but much less than in WT littermate embryos. *Ick* mutants show a much shorter femur and endochondral bone that display a significantly decreased proliferative zone (n=4).

In *Ick* mutant embryos, the vertebrae appears deformed and the IVD is defective (Fig. 4). Compared with WT, vertebrae of *Ick* mutants is much narrower, irregularly shaped, and severely distorted. A significant reduction of mineralization in the spinal column of *Ick* mutants is shown by Von Kossa staining (Fig. 5).

**Figure 4:**
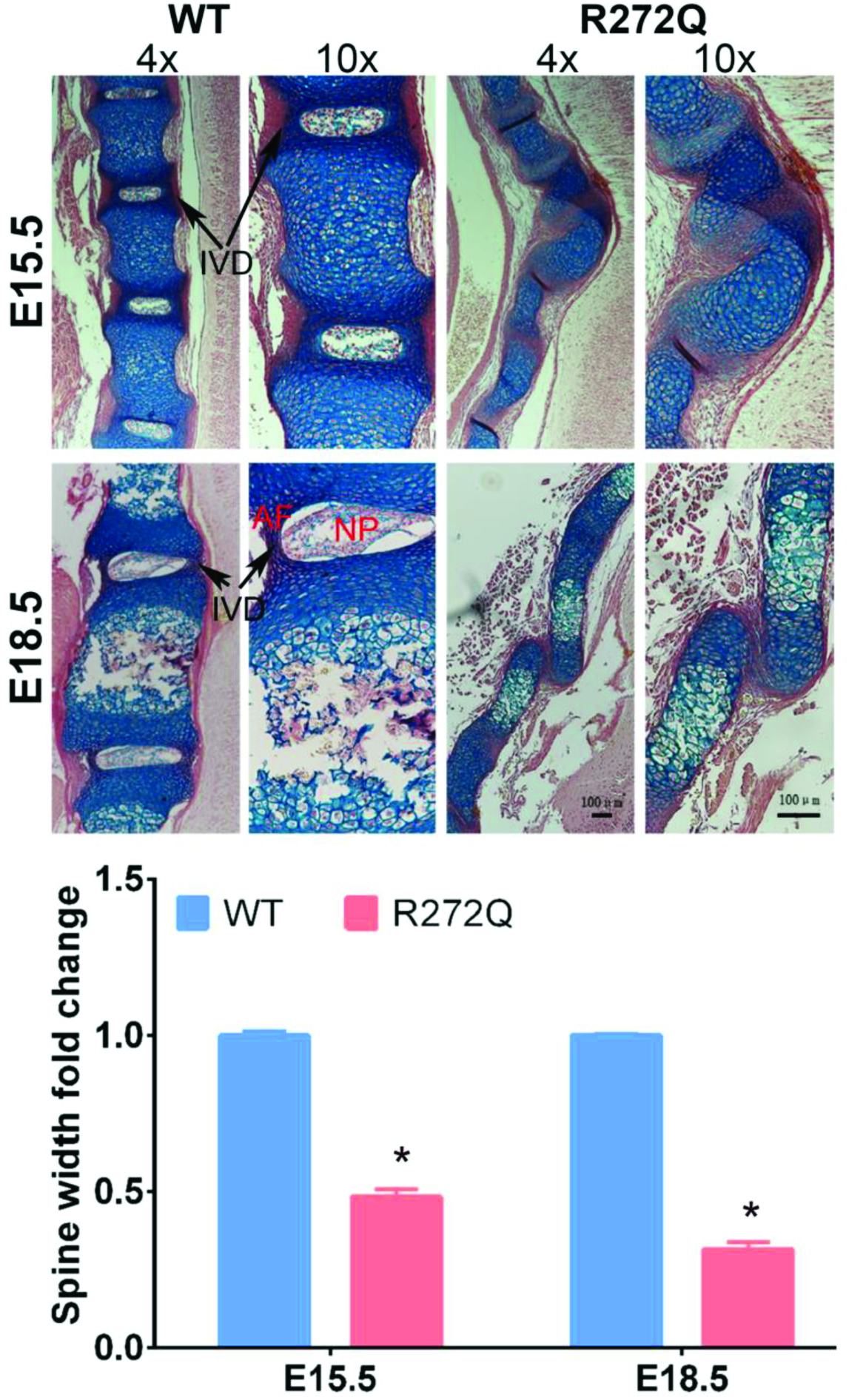
*Ick* mutant spine is distorted and deficient of intervertebral discs. In *Ick* mutant spine, only premature chondrocytes (E15.5) or hypertrophic chondrocytes (E18.5) locate in the spinal ossification center. The width of spinal column is much narrower in *Ick* mutants as compared with their WT littermates (n=3). Scale bar=100 µm.

**Figure 5:**
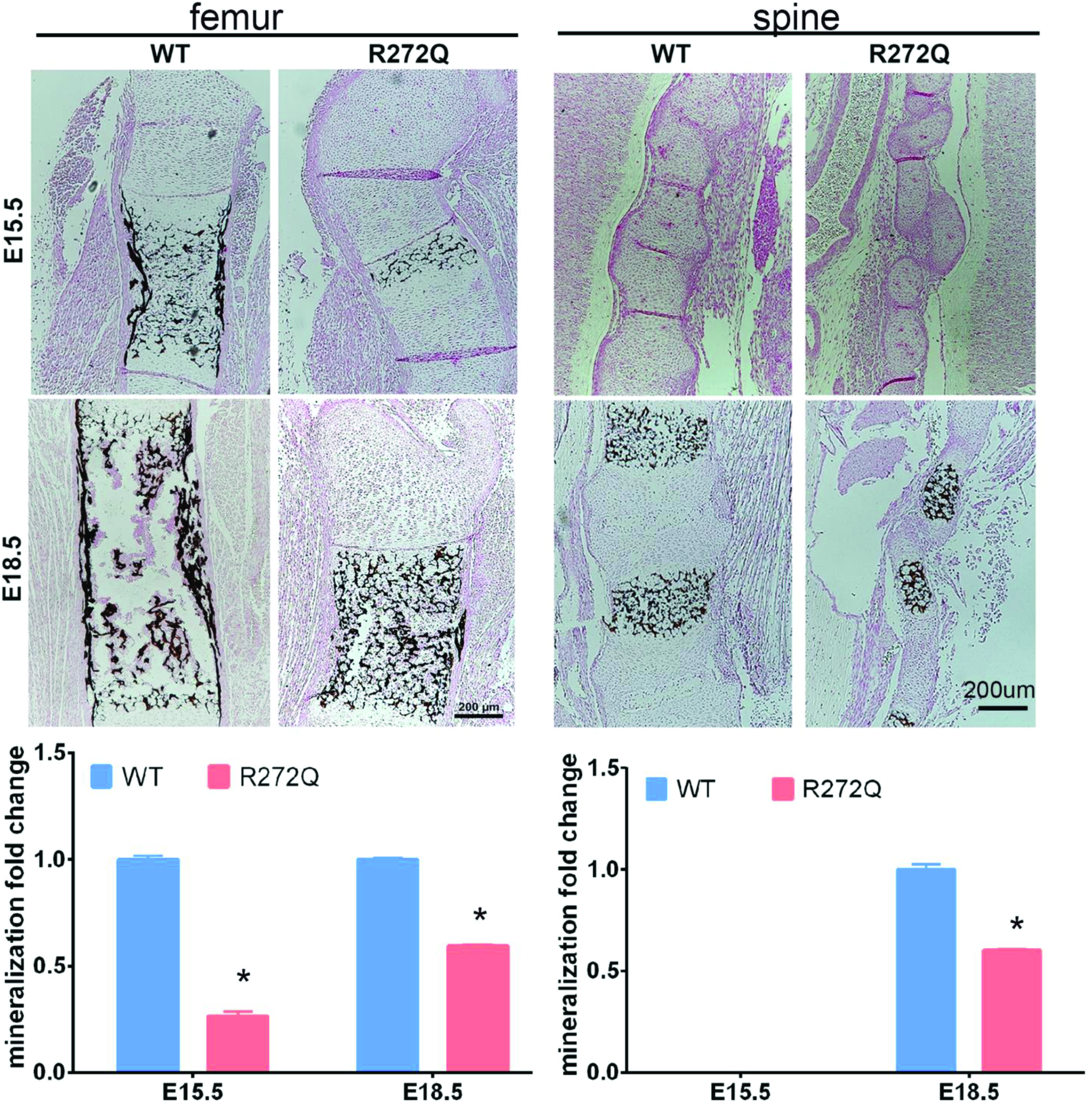
Mineralization of spinal column and femur is severely impaired in *Ick* mutant embryos. Paraffin embedded sections from E15.5 and E18.5 embryos stained by von Kossa showing mineralized bone in black. In E18.5 embryos, significantly less mineralization of femur and spine was observed in *Ick* mutant than in WT embryos (n=3).

### Abnormal chondrocytes maturation in *Ick* mutant skeletal system

Hypertrophic chondrocytes play an important role in bone mineralization. To determine whether impaired mineralization of *Ick* mutant embryos is due to compromised chondrocyte maturation, we performed immunostaining to compare the expression of collagen type II, a marker for chondrocytes. In E15.5 WT embryos, type II collagen is primarily expressed in the resting and proliferative chondrocytes and ECM in the growth plate of femurs (Fig. 6). It is also expressed in the pre-hypertrophic zone. In contrast, *Ick* mutant embryos express type II collagen throughout the proliferative region and into the hypertrophic region.

**Figure 6:**
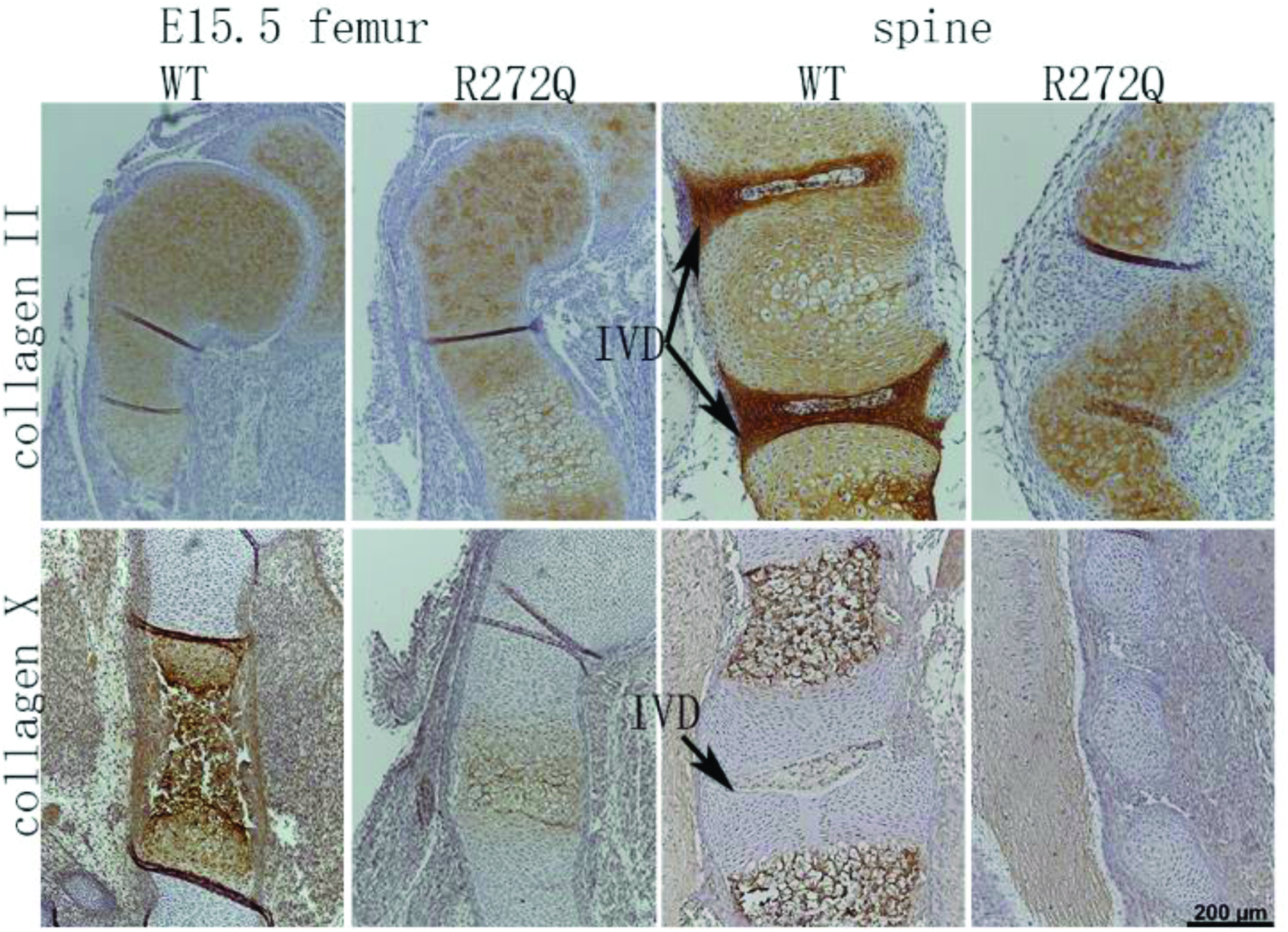
Loss of ICK functions significantly alters chondrocyte differentiation and maturation in spinal column and femur. Paraffin embedded sections from spinal columns and femurs of E15.5 WT and R272Q mutants were immunostained for collagen II or collagen X. Note that in the spinal and femur ossification centers of *Ick* mutant embryos, most of the cells were collagen II-positive chondrocytes and collagen X-positive terminally differentiated chondrocytes were barely detectable (spine) or markedly reduced (femur).

In WT spine, collagen type II expresses abundantly in annulus fibrosus (AF) tissue and proliferating chondrocytes (Fig. 6). In *Ick* mutants, IVDs are deficient, and collagen II expression was observed in proliferating and hypertrophic zones (Fig. 6). Type X collagen was observed in the hypertrophic regions and ECM of long bones and vertebrae in WT E15.5, but much less in *Ick* mutants. These results indicate that lack of ICK functions significantly impairs chondrocyte maturation in both spine and long bones.

### Reduced chondrocytes proliferation in *Ick* mutant skeletal system

Growth within the cartilage is dependent upon the proliferation of chondrocytes. To examine proliferation in *Ick* mutant embryos, we performed phospho-H3 staining of sections of the spine and long bones from *Ick* WT and mutant embryos. As shown in Figure 7, a marked reduction in the percentage of pH3 positive nuclei (brown color) in the growth plate of long bones and spines of *Ick* mutant embryos was observed as compared with WT. This data suggests that ICK is required to maintain high rate of chondrocyte proliferation in rapidly growing long bones and spinal columns during mouse skeletal development.

**Figure 7:**
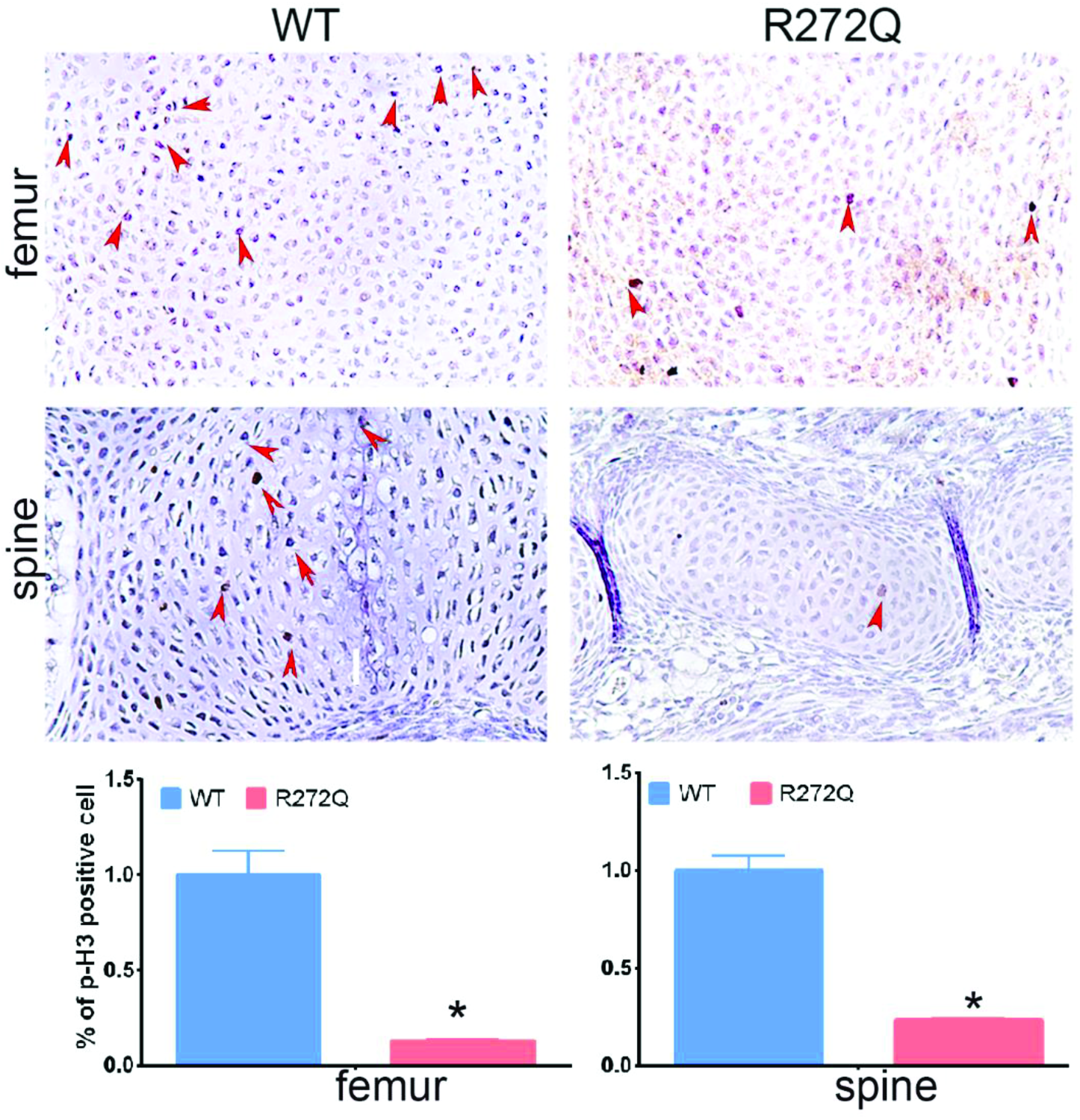
ICK dysfunction attenuates chondrocyte proliferation in spinal column and femur. Paraffin embedded sections from E15.5 of WT and *Ick* mutant embryos were immunostained with the proliferation marker p-Histone H3. Bar graphs show the quantitative data of positive cells (n=3). Scale bar=50 µm.

## Discussion

ECO is a rare recessive genetic disorders caused by a homozygous loss-of-function mutation R272Q in the human *ICK* gene. Recent studies from *Ick* knockout mouse models confirmed ECO phenotypes in the skeletal system; however, the cellular and molecular mechanisms underlying the skeletal defects in ECO syndrome are still elusive. Using an innovative ECO mutation knock-in mouse model, we hereby demonstrate that ICK functional deficiency not only reduces chondrocyte proliferation but also disrupts chondrocyte differentiation and maturation, resulting in severe deficiency in mineralization of long bones and spinal column.

*Ick*^R272Q/R272Q^ mutant revealed a marked disruption of growth plate architecture with a reduction in proliferative chondrocyte zone, consistent with what was observed earlier in *Ick*^−/−^ mutant [7, 9]. Here, we provide further evidence that a significant decrease in proliferating chondrocytes leads to the shortened PC zone of growth plate. Although the hypertrophic chondrocyte zone of *Ick*^R272Q/R272Q^ is not significantly different from that of WT, the number of hypertrophic chondrocytes expressing collagen X, a marker for terminally differentiated hypertrophic chondrocytes, is dramatically reduced concomitant with a severe deficiency in calcified cartilage matrix and a significant delay of mineralization in the long bone and spinal ossification center. In the final step of chondrocyte differentiation, the cartilaginous matrix is replaced by mineralized matrix [16]. Hypertrophic chondrocyte terminal differentiation is a critical step in mineralization. This developmental process requires stringent control mechanisms including action of hormones, morphogens, ECM proteins as well as transcriptional factors [17–18]. There are common factors that regulate both chondrocyte maturation and bone calcification [19]. *Ick* mutant embryos show a higher expression of collagen II in hypertrophic zone concomitant with a much lower type X collagen expression in the ossification center. These results provide new evidence for an essential role of ICK in chondrogenic cell differentiation and maturation during skeletal development.

A striking new skeletal phenotype of ECO syndrome identified in *Ick* R272Q mutant is the deficiency of IVD. IVD consists of gelatinous nucleus pulposus (NP) in the center, surrounded by fibrocartilaginous annulus fibrosus (AF), and superior and inferior cartilaginous endplates. NP is believed to be derived from the notochord while AF and endplate are developed from sclerotome [20]. Notochord formation is the foundation of the disc development because notochord is a crucial signaling center for the paraxial mesoderm that gives rise to the sclerotome and subsequently fibrous AF tissue and cartilage endplate of IVD [21–22]. NP is apparent by E15.5 as shown in Fig. 4, composed of notochordal cells and large vacuolated cells. Disruption of notochord formation and/or sclerotome specification can lead to a much smaller or completely absence of IVD. Thus, the absence of IVD in *Ick* mutant spine strongly implicates an essential role for ICK in the determination of early notochord and sclerotome formation which requires further investigation in our future studies.

In summary, we demonstrate that *Ick* R272Q knock-in mouse model resembles the clinical features of ECO, and ICK plays an important role in bone and IVD development by regulating chondrocyte proliferation, differentiation and maturation.

## Acknowledgments

We are grateful to financial support partially from NIH R01AR064792 (XL), DK082614 and CA195273 (ZF). We appreciate the technical assistance of the Research Histology Core at University of Virginia. The funders have no role in the study.

---

**Supplemental Figure 1:** ICK expression is detected in the bone and cartilage tissue. (**A**) Western blot showing ICK expression in tibia, sternum, and rib. (**B**) LacZ staining showing ICK expression in the spine and limbs. Scale bar=200 µm

Supplemental Method: *In-situ* Lac-Z staining

The heterozygous *Ick-LacZ* knockout/reporter mice (strain ID: *Ick^tm1a(KOMP)Mbp^*) were obtained from the Knockout Mouse Project (KOMP) Repository. Whole embryos were fixed in 4% paraformaldehyde (PFA) solution for two hours, followed by rinses in PBS containing 0.01% sodium deoxycholate and 0.02% NP-40 before incubation with X-gal (1 mg/ml) solution at RT until color developed to desired intensity.

